# Genomic analysis provides novel insights into diversification and taxonomy of *Allorhizobium vitis* (i.e. *Agrobacterium vitis*)

**DOI:** 10.1101/2020.12.19.423612

**Authors:** Nemanja Kuzmanović, Enrico Biondi, Jörg Overmann, Joanna Puławska, Susanne Verbarg, Kornelia Smalla, Florent Lassalle

## Abstract

**Background:** *Allorhizobium vitis* (formerly named *Agrobacterium vitis* or *Agrobacterium* biovar 3) is the primary causative agent of crown gall disease of grapevine worldwide. We obtained and analyzed whole-genome sequences of diverse *All. vitis* strains to get insights into their diversification and taxonomy.

**Results:** Pairwise genome comparisons and phylogenomic analysis of various *All. vitis* strains clearly indicated that *All. vitis* is not a single species, but represents a species complex composed of several genomic species. Thus, we emended the description of *All. vitis*, which now refers to a restricted group of strains within the *All. vitis* species complex (i.e. *All. vitis sensu stricto*) and proposed a description of a novel species, *All. ampelinum* sp. nov. The type strain of *All. vitis sensu stricto* remains the current type strain of *All. vitis*, K309^T^. The type strain of *All. ampelinum* sp. nov. is S4^T^. We also identified sets of gene clusters specific to the *All. vitis* species complex, *All. vitis sensu stricto* and *All. ampelinum*, respectively, for which we predicted the biological function and infer the role in ecological diversification of these clades, including some we could experimentally validate. *All. vitis* species complex-specific genes confer tolerance to different stresses, including exposure to aromatic compounds. Similarly, *All. vitis sensu stricto*-specific genes confer the ability to degrade 4-hydroxyphenylacetate and a putative compound related to gentisic acid. *All. ampelinum*-specific genes have putative functions related to polyamine metabolism and nickel assimilation. Congruently with the genome-based classification, *All. vitis sensu stricto* and *All. ampelinum* were clearly delineated by MALDI-TOF MS analysis. Moreover, our genome-based analysis indicated that *Allorhizobium* is clearly separated from other genera of the family *Rhizobiaceae*.

**Conclusions:** Comparative genomics and phylogenomic analysis provided novel insights into the diversification and taxonomy of *Allorhizobium vitis* species complex, supporting our redefinition of *All. vitis sensu stricto* and description of *All. ampelinum*. Our pan-genome analyses suggest that these species have differentiated ecologies, each relying on specialized nutrient consumption or toxic compound degradation to adapt to their respective niche.

## Background

*Allorhizobium vitis* (formerly named *Agrobacterium vitis* or *Agrobacterium* biovar 3) is a bacterium primarily known as a plant pathogen causing crown gall disease of grapevine (*Vitis vinifera*) (1). This economically important plant disease may cause serious losses in nurseries and vineyards. *All. vitis* is widely distributed pathogen, detected in almost all grapevine growing regions throughout the world. This bacterium seems to be associated almost exclusively with grapevine. It has been isolated from crown gall tumors, xylem sap, roots, rhizosphere, non-rhizosphere soil of infected vineyards, decaying grape roots and canes in soil, but also from the phyllosphere of grapevine plants (reviewed in (1)). In one exceptional case, *All. vitis* was isolated from galls on the roots of kiwi in Japan (2).

*All. vitis* is an aerobic, non-spore-forming, Gram-negative, rod-shaped bacterium with peritrichous flagella (3). It is a member of the alphaproteobacterial family *Rhizobiaceae*, together with other genera hosting tumor-inducing plant pathogens, including *Agrobacterium* and *Rhizobium*. With time, the taxonomy of *All. vitis* has undergone various changes. Tumorigenic strains associated with crown gall of grapevine were initially defined as an atypical group that could neither be classified as *Agrobacterium* biovar 1 (i.e., *Agrobacterium tumefaciens* species complex) nor as biovar 2 (i.e. *Rhizobium rhizogenes*) (4). Afterwards, several studies classified these atypical strains as *Agrobacterium* biovar 3 (biotype 3), based on their biochemical and physiological characteristics (5-7). Serological analysis using monoclonal antibodies also allowed differentiation of *Agrobacterium* biovar 3 strains (8). Polyphasic characterization involving DNA-DNA hybridization (DDH), phenotypic and serological tests clearly showed that *Agrobacterium* biovar 3 strains represent a separate species, for which the name *Agrobacterium vitis* was proposed (9). However, multi-locus sequence analysis (MLSA) suggested that *A. vitis* is phylogenetically distinct from the genus *Agrobacterium*, and prompted the transfer of this species to the revived genus *Allorhizobium* (10, 11).

The genus *Allorhizobium* was created by de Lajudie et al. (12) and initially included single species *Allorhizobium undicola*. Afterwards, Young et al. (13) proposed reclassification of *All. undicola* and its inclusion into the genus *Rhizobium*, while Costechareyre et al. (14) suggested that this species might belong to the genus *Agrobacterium*. However, these studies employed single gene phylogenies, which were insufficient to support such taxonomic revisions. The authenticity of the genus *Allorhizobium* and the clustering of *All. vitis* within it was unequivocally confirmed by genome-wide phylogenies (15, 16). Moreover, distinctiveness of *All. vitis* with respect to the genus *Agrobacterium* was further supported by their different genome organization, with the genus *Agrobacterium* being characterized by the presence of a circular chromosome and a secondary linear chromid (17, 18). Chromids are defined as large non-dispensable plasmids carrying essential functions (19). In contrast to *Agrobacterium*, the *All. vitis* strains carry two circular chromosomes (18, 20, 21). However, the smaller circular chromosome (named chromosome II) was later classified as a chromid in the fully sequenced strain *All. vitis* S4^T^ (19). Additionally, genomes of *All. vitis* and other agrobacteria include a variable number of plasmids.

In recent years, genomics has significantly impacted the taxonomy of bacteria, leading to the revisions in classification of different bacterial taxa. In particular, a novel genomics-based taxonomy primarily relies on the calculation of various overall genome relatedness indices (OGRIs) and estimation of genome-based phylogenies (22-24), largely replacing the traditionally used methods of 16S rRNA gene phylogeny and DDH (25, 26). Genomic information were also highly recommended as essential for the description of new rhizobial and agrobacterial taxa (27). In addition, it has been recommended that some functions and phenotypic characters may not be considered for taxonomic classification. This particularly applies to the tumor-inducing ability of agrobacteria, which is mainly associated with the dispensable tumor-inducing (Ti) plasmid.

Information on genetic diversity and relatedness of strains responsible for crown gall disease outbreaks provide important insights into the epidemiology, ecology and evolution of the pathogen. Numerous studies indicated that *All. vitis* strains are genetically very diverse (reviewed in (1)). In our previous study, we analyzed a representative collection of *All. vitis* strains originating from several European countries, Africa, North America, and Australia using MLSA, which indicated a high genetic diversity between strains, clustered into four main phylogenetic groups (28). These data suggested that *All. vitis* might not be a homogenous species, but a species complex comprising several genomic species, warranting further investigation of the diversification and evolution of *All. vitis* towards a more complete elucidation of its taxonomy.

In this work, we selected representative strains belonging predominantly to the two most frequent phylogenetic groups identified in our previous study (28) that included the well-studied *All. vitis* type strain K309^T^ and the fully sequenced strain S4^T^, respectively. We obtained draft genome sequences for 11 additional strains and performed comparative genomic and phylogenetic analyses to reveal the diversification history and synapomorphies of these groups. In parallel, we investigated phenotypic features of selected strains. The combination of these approaches allows us to revise the taxonomy within this group, notably by emending the description of *All. vitis* (*All. vitis sensu stricto*) and proposing the new species *All. ampelinum*.

## Results

### *Allorhizobium vitis* genome sequencing

Draft genome sequences were obtained for 11 *All. vitis* strains (Table 1), with average coverage depth ranging from 65- to 96-fold. The total size of draft genome assemblies ranged from 5.67 to 6.52 Mb, with a GC content ranging 57.5-57.6% (Table 1), which was similar to the genomes of other *All. vitis* strains sequenced so far (Table S1b).

**Table 1.**
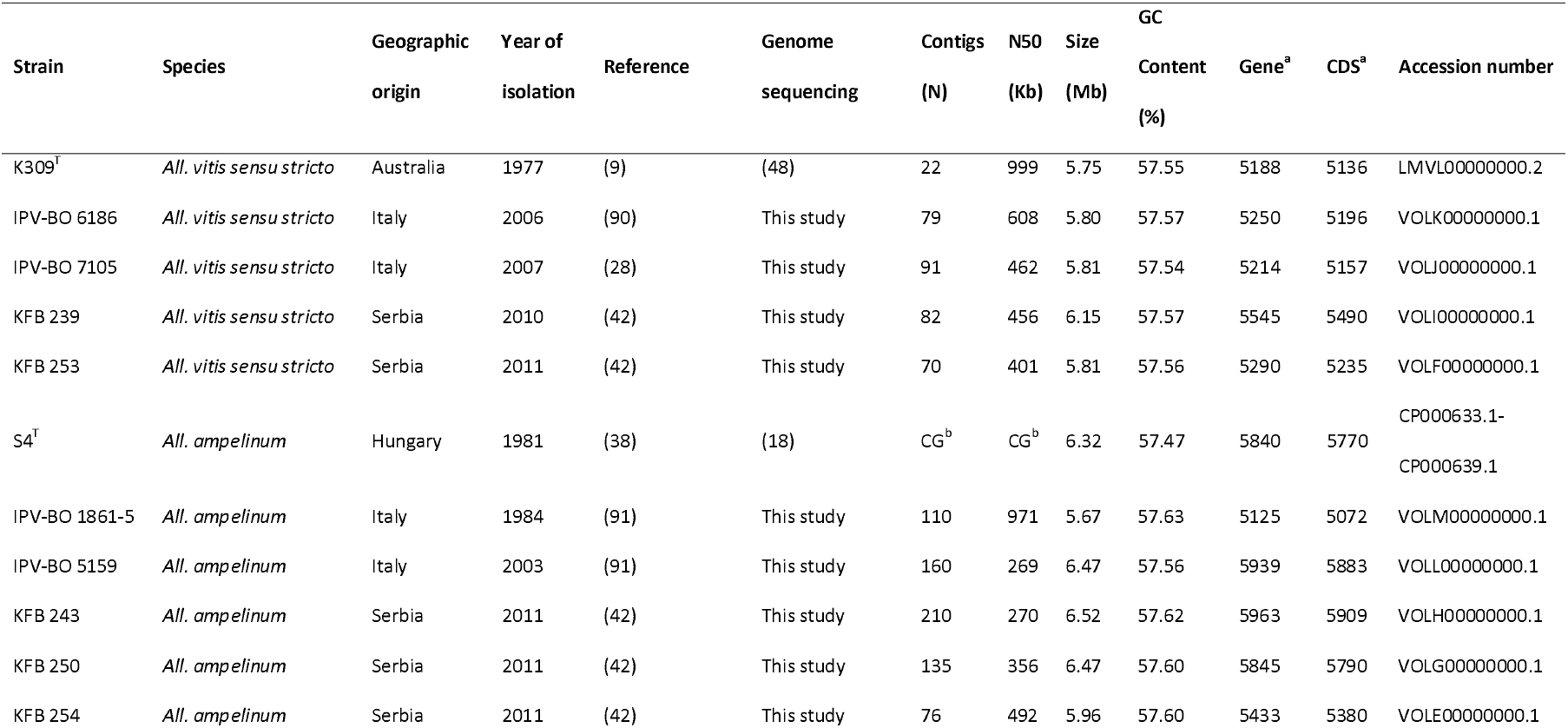
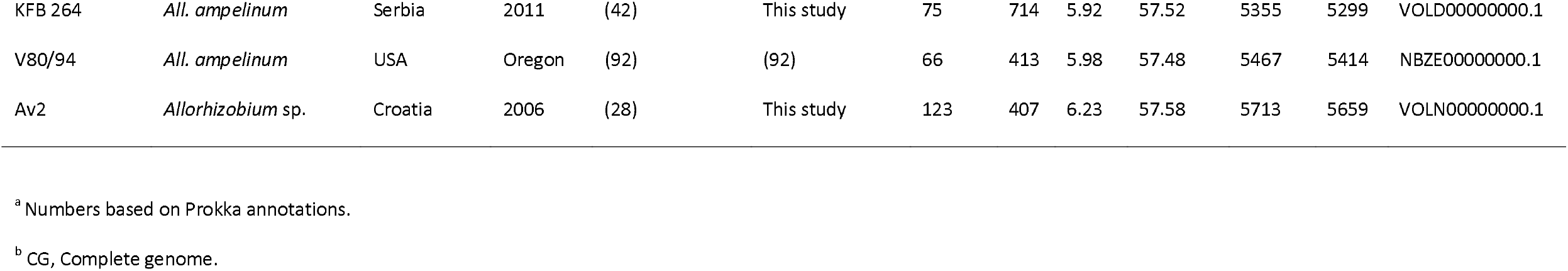
Characteristics of 14 strains of the *All. vitis* species complex analyzed in this study and their genome sequence features

### Core-genome phylogeny and overall genome relatedness indexes measurements

A core-genome phylogeny was inferred for 14 strains of *All. vitis* (Table 1) and 55 reference *Rhizobiaceae* strains (Table S1a). A phylogenomic tree that was reconstructed from the concatenation of 344 non-recombining core marker genes confirmed the grouping of *Allorhizobium* species separately from other *Rhizobiaceae* genera (Figs. 1 and S1). The clade comprising all members of the genus *Allorhizobium* was well separated from its sister clade, which included members of the group provisionally named “*R. aggregatum* complex” (11), as well as representatives of the genus *Ciceribacter*.

**Fig. 1.**
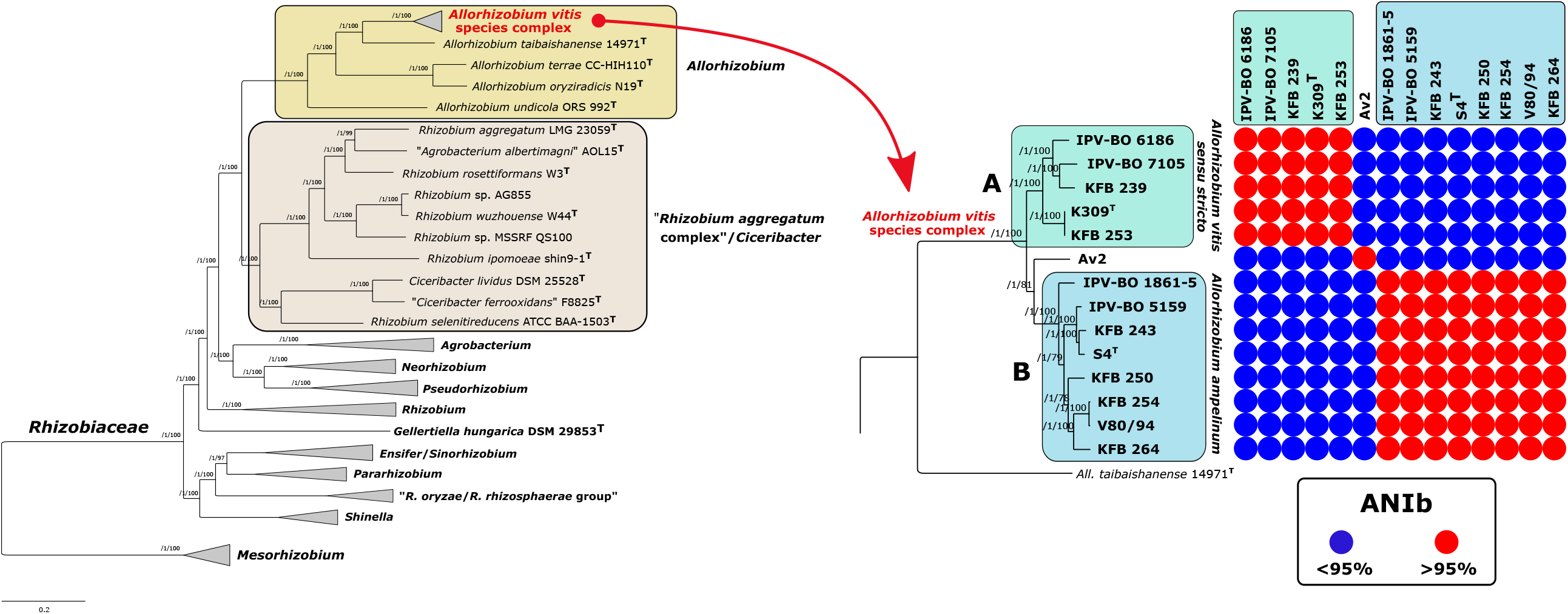
Maximum-likelihood core-genome phylogeny of 69 strains belonging to the genus *Allorhizobium* and other *Rhizobiaceae* members (part collapsed). The tree was estimated with IQ-TREE from the concatenated alignment of 344 top-ranked genes selected using GET_PHYLOMARKERS software. The numbers on the nodes indicate the approximate Bayesian posterior probabilities support values (first value) and ultra-fast bootstrap values (second value), as implemented in IQ-TREE. The tree was rooted using the *Mesorhizobium* spp. sequences as the outgroup. The scale bar represents the number of expected substitutions per site under the best-fitting GTR+F+ASC+R6 model. “*Allorhizobium vitis*” clade is collapsed on the left tree and shown expanded on the right. The matrix represents the distribution of ANIb values for the genomic sequences of the clade corresponding to the *All. vitis* species complex, relative to the typical species delimitation threshold of 95%. The same tree, but without collapsing clades, is presented in the Figure S1.

*All. vitis* strains formed a well-delineated clade within the *Allorhizobium* genus (Figs. 1 and S1). Furthermore, *All. vitis* strains were clearly differentiated into two well-supported sub-clades (clades A and B), while strain Av2 branched separately from each of these two clades (Figs. 1 and S1). OGRIs values (Table S2) indicated that sub-clades A and B, as well as strain Av2, represent separate genomic species. In other words, the core-genome phylogeny and OGRI measurements showed that *All. vitis* is not a single species, but a species complex composed of at least three separate genomic species.

The first genomic species, corresponding to sub-clade A, comprises the type strain of *All. vitis* (strain K309^T^) (Fig. 1). Although dDDH values suggested that the cluster containing strains K309^T^ and KFB 253 might belong to a separate species compared to other strains comprised in this sub-clade (Table S2e), this was not supported by the other four OGRIs calculated here (Table S2a-d). Indeed, dDDH values for these strains (65.9-66.4 %) were relatively close to the generally accepted threshold value of 70 %. A revised description of the species *All. vitis*, hereafter referred to as *All. vitis sensu stricto*, is given below.

The second genomic species, corresponding to sub-clade B, included eight strains originating from various geographic areas (Table 1; Fig. 1). It included the well-studied strain S4^T^, whose high-quality genome sequence was described previously (18). The dDDH value obtained from the comparison of strain KFB 254 with strain IPV-BO 1861-5 was below, but very close to the 70 % threshold value generally accepted for species delineation (Table S2e). However, other OGRIs unanimously indicated that strains from this sub-clade belong to the same species (Table S2a-d). A description of the novel species corresponding to sub-clade B, for which the name *Allorhizobium ampelinum* sp. nov. is proposed, is given below.

The third genomic species comprised strain Av2 alone (Figs. 1 and S1, Table S2). To get a more comprehensive insight into the diversity of the *All. vitis* species complex, we conducted a second phylogenomic analysis where we included 34 additional genomes of *All. vitis* that were available in GenBank but not yet published (Table S1b). Based on core-genome phylogeny and ANIb calculations (Fig. S2, Table S3), additional strains were taxonomically assigned as *All. vitis sensu stricto* (sub-clade A) and *All. ampelinum* (sub-clade B). Strain Av2 then grouped with three other strains originating from the USA (sub-clade D; Fig. S2). These four strains comprised in the sub-clade D were genetically very similar and exhibited >99.8 ANI between each other (Table S3). Moreover, additional sub-clades C and E were apparent, corresponding to two other new genomic species of *All. vitis* species complex (Fig. S2, Table S3). Genomic species corresponding to sub-clade C and sub-clade D were closely related, as their ANIb values were in the range 94.62-94.93 %, which is slightly below the threshold for species delimitation (∼95-96%) (29).

### Pan-genome analyses

A ML pan-genome phylogeny of the 64 *Rhizobiaceae* genome dataset was estimated from a matrix of the presence or absence of 33,396 orthologous gene clusters (Fig. 2; Fig. S3). The pan-genome phylogeny (Fig. 2; Fig. S3) presented the same resolved sub-clades of the *All. vitis* complex as the core-genome phylogeny (Fig. 1). Furthermore, *Rhizobiaceae* genera and clades were generally differentiated based on the pan-genome tree (Fig. 2; Fig. S3). Nevertheless, some inconsistencies were observed: tumorigenic strain *Neorhizobium* sp. NCHU2750 was more closely related to the representatives of the genus *Agrobacterium*, while nodulating *Pararhizobium giardinii* H152^T^ was grouped with *Ensifer* spp. (Fig. 2; Fig. S3). These inconsistencies were also observed in another pan-genome phylogeny inferred using parsimony (data not shown). Such limitations of gene content-based phylogenies have previously been reported (30, 31).

**Fig. 2.**
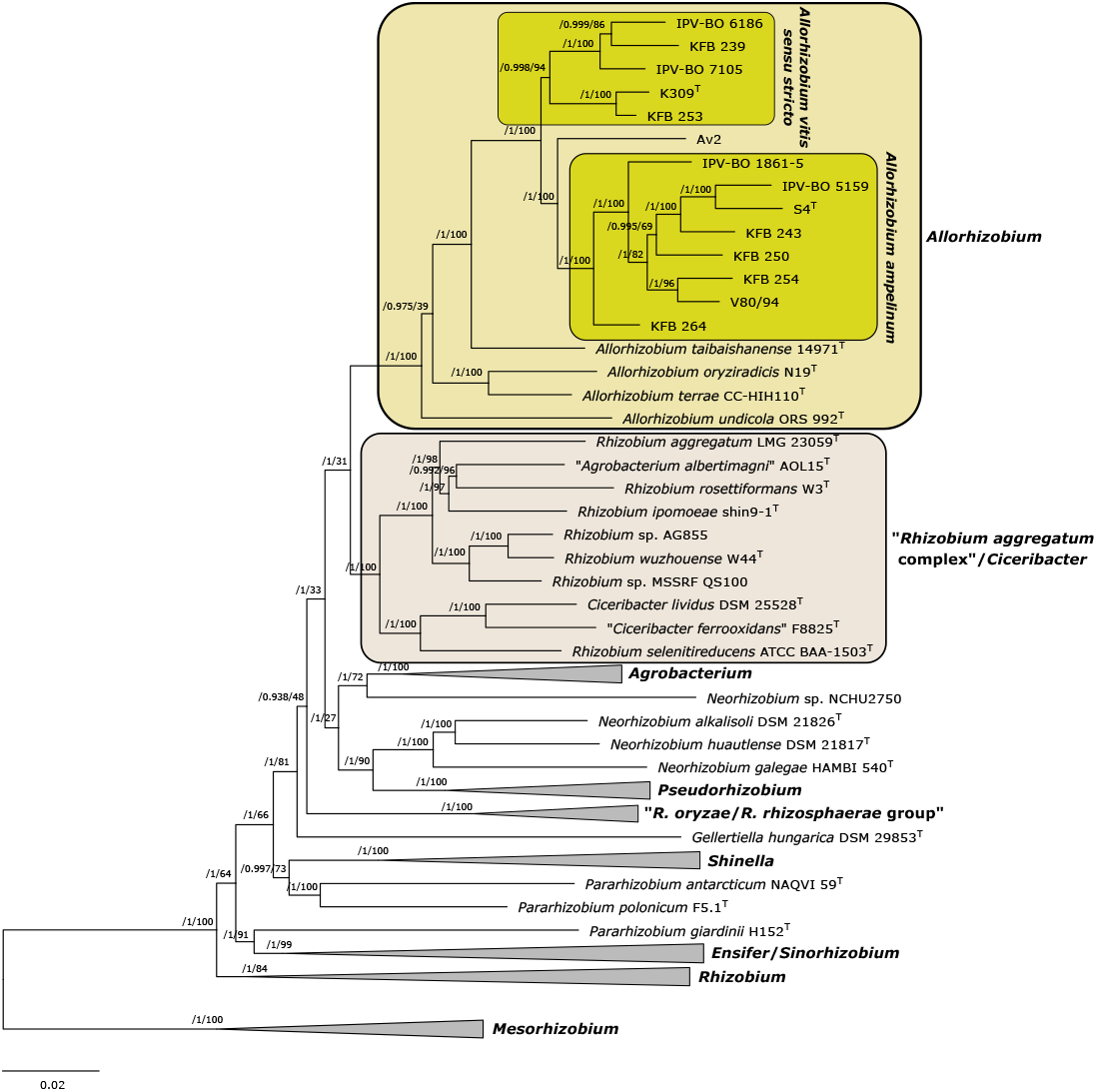
Maximum-likelihood pan-genome phylogeny of 69 strains belonging to the genus *Allorhizobium* and other *Rhizobiaceae* members (part collapsed). The tree was estimated with IQ-TREE from the consensus (COGtriangles and OMCL clusters) gene presence/absence matrix containing 33,396 clusters obtained using GET_HOMOLOGUES software. The numbers on the nodes indicate the approximate Bayesian posterior probabilities support values (first value) and ultra-fast bootstrap values (second value), as implemented in IQ-TREE. The tree was rooted using the *Mesorhizobium* spp. sequences as the outgroup. The scale bar represents the number of expected substitutions per site under the best-fitting GTR2+FO+R5 model. The same tree, but without collapsing clades, is presented in the Figure S3.

Focusing on 14 *All. vitis* species complex strains, we identified 10,501 pan-genome gene clusters. The core-genome (‘strict core’ and ‘soft core’ compartments) of the species complex comprised 3,775 gene clusters (35.95% of total gene clusters), with 3,548 gene clusters strictly present in all 14 strains (Fig. 3). The accessory genome contained 4,516 in the cloud (43% of total gene clusters) and 2,210 gene clusters in the shell (21.05% of total gene clusters) (Fig. 3).

**Fig. 3.**
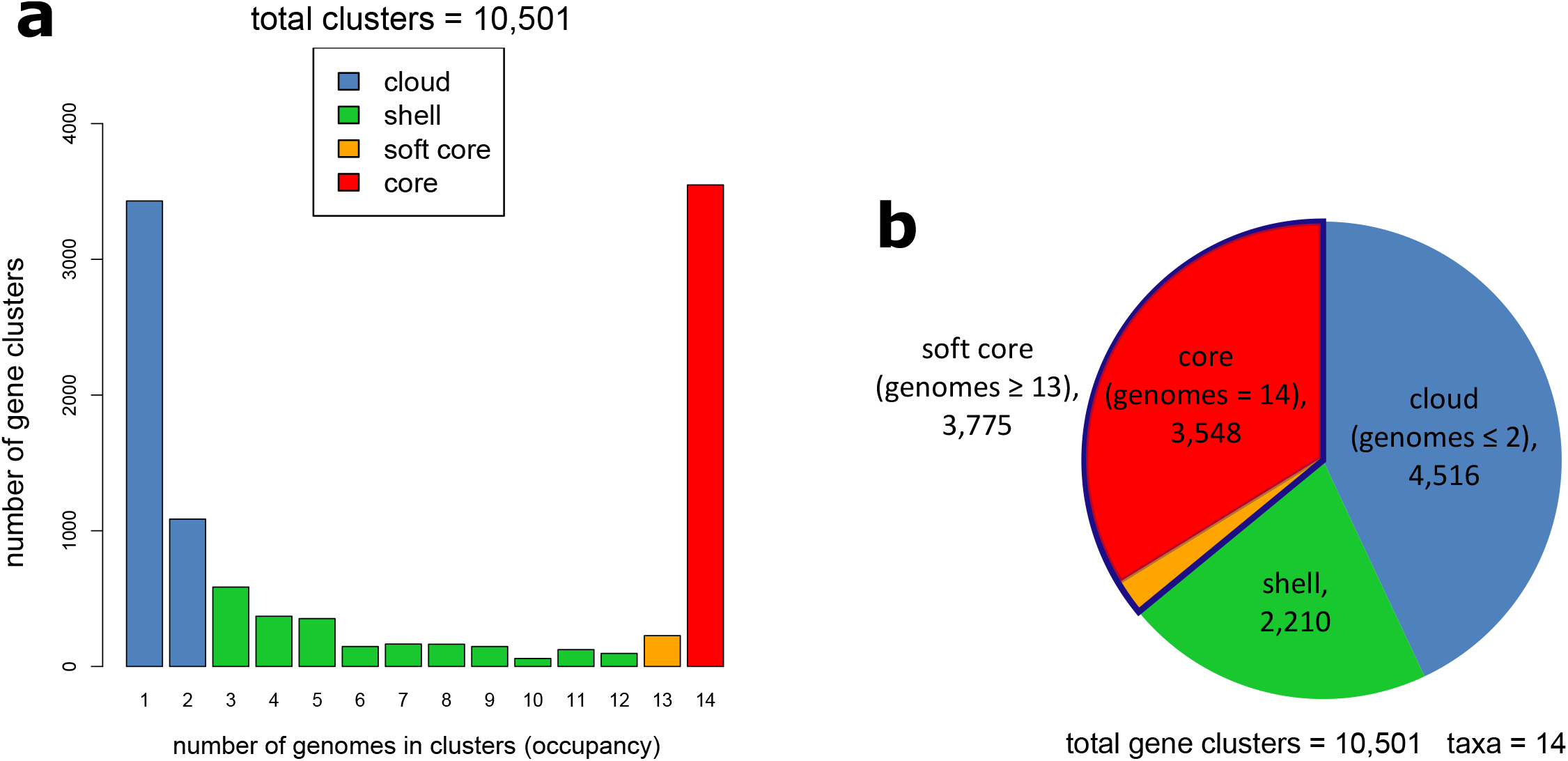
Pan-genome analyses of 14 *All. vitis* species complex strains. **a)** Bar plot showing the frequencies of orthologous clusters as predicted by the COGtriangles and OMCL algorithms. **b)** Pie chart showing the relative sizes (cluster numbers) contained in the core, soft-core, shell, and cloud genome compartments.

### Clade-specific gene clusters

Homologous gene families specific to particular clades of interest, i.e. with contrasted presence pattern with respect to closely related clades, were identified using both Pantagruel or GET_HOMOLOGUES software packages. Both sets of inferred clade-specific genes were to a large extent congruent, although some differences were observed (Table S4), owing to the distinct approaches employed by these software packages (32, 33). We focused on clusters of contiguous clade-specific genes for which we could predict putative molecular functions or association to a biological process. The results are summarized below and in Table S4.

#### *All. vitis* species complex

Based on Pantagruel and GET_HOMOLOGUES analyses, we identified 206 and 236 genes, respectively, that are specific to the *All. vitis* species complex (*Av*SC-specific genes), i.e. present in all strains of *All. vitis sensu stricto, All. ampelinum* and *Allorhizobium* sp. Av2, and in no other *Allorhizobium* strain. *Av*SC-specific genes are mostly located on the second chromosome (chromid). While some *Av*SC-specific genes are found on the Ti plasmid and include the type 4 secretion system, this likely only reflects a sampling bias whereby all *All. vitis* species complex strains in our sample were tumorigenic and possessed a Ti plasmid. As such, Ti plasmid-encoded genes directly associated with pathogenicity were not further considered or discussed in this study.

Half of the *Av*SC-specific genes are gathered in contiguous clusters for most of which we could predict putative function (Table S4); most of the other half are scattered on chromosome 1 and have unknown function. Predicted functions of clustered genes revealed that they are strikingly convergent: most are involved in either environmental signal perception (4 clusters), stress response (2 clusters), aromatic compound and secondary metabolite biosynthesis (3 clusters) and/or aromatic compound degradation response (2 clusters). In addition, one cluster encodes a multicomponent K^+^:H^+^ antiporter, which is likely useful for adaptation to pH changes, and three clusters harbor several ABC transporter systems for sugar or nucleotide uptake. Finally, one cluster on chromosome 1 encodes a putative auto-transporter adhesin protein, which may have a role in plant commensalism and pathogenesis.

All studied *All. vitis* species complex strains carried a *pehA* gene encoding a polygalacturonase enzyme. Unlike other agrobacteria, *All. vitis* strains are known to produce a polygalacturonase, regardless of their tumorigenicity (34). However, this gene was present also in *All. taibaishanense* 14971^T^, *All. terrae* CC-HIH110^T^ and *All. oryziradicis* N19^T^, but absent in *All. undicola* ORS 992^T^ and in other studied members of the *Rhizobiaceae* family.

Furthermore, we detected the presence of gene encoding enzyme 1-aminocyclopropane-1-carboxylate deaminase (*acdS*) in all studied *All. vitis* species complex strains. This gene is considered to be important for plant-bacteria interaction through its involvement in lowering the level of ethylene produced by the plant (35). We found this gene in all other *Allorhizobium* spp., and in some other *Rhizobiaceae* (data not shown), including *R. rhizogenes* strains. However, *acdS* gene was not present in *Agrobacterium* spp., even when the similarity search (blastp) was extended to *Agrobacterium* spp. strains available in GenBank, consistent with previous findings (36).

Tartrate utilization ability was previously reported for most of the *All. vitis* strains (37-39). Therefore, we searched *All. vitis* species complex genomes for the presence of tartrate utilization (TAR) regions. All strains except IPV-BO 6186 and IPV-BO 7105 carried TAR gene clusters. Moreover, we could not find any *All. vitis*-like TAR regions in any other *Rhizobiaceae* strain. Sequence comparison of TAR regions from *All. vitis* species complex strains using ANIb algorithm (Table S5) showed they could be divided into four types (Fig. S4). The first type is represented by a previously characterized TAR region called TAR-I, carried on the tartrate utilization plasmid pTrAB3 of strain AB3 (39, 40). The second type included representatives of TAR-II (carried on pTiAB3) and TAR-III (carried on pTrAB4) regions, which were previously described to be related to each other (39, 41). A third TAR region type, which we designate TAR-IV, was characterized by the absence of a second copy of *ttuC* gene (tartrate dehydrogenase). The TAR-IV region type is found in *All. ampelinum* strain S4^T^, in which the tartrate utilization system is located on the large plasmid pAtS4c (initially named pTrS4) (40). The TAR system of *Allorhizobium* sp. strain Av2 is a unique type (TAR-V), which is related to region type TAR-I, but is characterized by the absence of the *ttuA* gene (a LysR-like regulator). We compared the distribution of these TAR region types in strain genomes, showing there is no TAR region type associated to any genomic species (Table S6). *All. vitis sensu stricto* strains K309^T^ and KFB 253 carry a TAR-II/III region. In addition to TAR-II/III region, strain KFB 239 carries a TAR-I region (Table S6), a combination similar to that found in the well-characterized strain AB3 (39). *All. ampelinum* strains S4^T^, IPV-BO 1861-5, KFB 264 and V80/94 contain a TAR-IV region, while the remaining *All. ampelinum* strains IPV-BO 5159, KFB 243, KFB 250 and KFB 254 additionally carry a TAR-II/III region (Table S6).

#### All. vitis sensu stricto

Using Pantagruel and GET_HOMOLOGUES pipelines, we identified 63 and 78 genes, that are specific to *All. vitis sensu stricto* (Av-specific, present in all five strains and in none of *All. ampelinum*), respectively. 32 of these Av-specific genes are clustered into four main loci in the genome of strain K309^T^, for which we could predict putative function (Table S4). One *Av*-specific gene cluster (Av-GC1, Table S4) comprised genes functionally annotated to be involved in the degradation process of salicylic acid and gentisic acid (2,5-dihydroxybenzoic acid) (MetaCyc pathways PWY-6640 and PWY-6223). Av-GC1 was located on Contig 1 (LMVL02000001.1) of reference strain K309^T^ genome, which is likely part of the chromid, based on its high ANI with the chromid (Chromosome 2) of strain S4^T^, whose genome sequence is complete. BLAST searches showed that this gene cluster is also present in some representatives of *Agrobacterium deltaense*, i.e. *Agrobacterium* genomospecies G7 (data not shown). Av-GC1 is predicted to encode the degradation of salicyl-CoA, an intermediate in degradation of salicylic acid, to 3-fumarylpyruvate, via gentisic acid. Interestingly, strains KFB 239, IPV-BO 6186 and IPV-BO 7105 carried additional genes encoding the degradation of salicylaldehyde to salicyl-CoA via salicylic acid and salicyl adenylate, as well as the gene encoding the final step of gentisic acid degradation, the conversion of 3-fumarylpyruvate to fumarate and pyruvate. The three strains encoding enzymes of the complete pathway for degradation of salicylic acid and gentisic acid, and remaining strains K309^T^ and KFB 253 carrying a partial gene cluster, were phylogenetically separated and formed distinct sub-clades within *All. vitis sensu stricto* (Fig. 1).

Another *Av*-specific gene cluster (Av-GC4, Table S4) was annotated to be involved in the degradation of 4-hydroxyphenylacetate (MetaCyc pathway 3-HYDROXYPHENYLACETATE-DEGRADATION-PWY). Gene content and comparative analysis of the contig carrying this gene cluster suggested that Av-GC4 is carried on a putative plasmid of *All. vitis sensu stricto* (data not shown).

In addition, *Av*-specific gene clusters Av-GC2 and Av-GC3 (Table S4) were both predicted to be involved in amino-acid uptake and catabolism. However, we were not able to predict the precise molecular function of proteins and substrates of enzymes encoded by these *Av*-specific gene clusters. Both these gene clusters are likely located on a putative plasmid, as suggested by the presence of plasmid-related genes (replication and/or conjugation associated genes) on the same contigs.

#### All. ampelinum

Based on Pantagruel and GET_HOMOLOGUES analyses, we identified 97 genes and 128 genes, respectively, that are specific to *All. ampelinum* (*Aa*-specific, present in all 8 strains and in none of *All. vitis sensu stricto*). Taking advantage of the finished status of strain S4^T^ genome, we found that 52/97 specific genes identified by Pantagruel occur on plasmids rather than chromosomes. This is a significant over-representation compared to the distribution of all genes (21.4% on plasmids, Chi-squared test *p*-value < 10^−6^) or core-genome genes (5.8% on plasmids, Chi-squared test *p*-value < 10^−16^). For eleven contiguous gene clusters we could predict putative function (Table S4). The *Aa*-specific gene clusters encode a variety of putative biological functions; an enrichment analysis of their functional annotations revealed a set of high-level biological processes that were over-represented: transport and metabolism of amino-acids or polyamines like putrescine (three separate clusters), lysin biosynthesis (two separate clusters), and nickel assimilation. The latter function is predicted for gene cluster Aa-GC10, which is located on the 631-kb megaplasmid pAtS4e and encodes the NikABCDE Ni^2+^ import system and a nickel-responsive transcriptional regulator NikR. Aa-GC10 additionally includes genes with predicted functions such as cation-binding proteins and a chaperone/thioredoxin, which may be involved in the biosynthesis of ion-associated cofactors.

### Phenotypic and MALDI-TOF MS characterization

The phenotypic properties of the newly described species *All. ampelinum* are listed in Table 2. API 20NE and Biolog GEN III analyses did not reveal clear discriminative features between *All. vitis sensu stricto* and *All. ampelinum*. However, a weak positive reaction for 4-hydroxyphenylacetic (p-hydroxy-phenylacetic) acid for strains belonging to *All. vitis sensu stricto* was recorded, unlike for those belonging to *All. ampelinum*, which were clearly negative. As bioinformatic analyses suggested that *All. vitis sensu stricto* strains carry a gene cluster encoding the degradation of 4-hydroxyphenylacetate, the metabolism of this compound was assayed in a separate biochemical test. Our results indicated that all *All. vitis sensu stricto* strains tested are able to metabolize 4-hydroxyphenylacetate, which was recorded by a vigorous bacterial growth and a change of pH (∼7.2 to ∼6.5), indicating the production of acid from the substrate oxidation. On the other hand, *All. ampelinum* strains showed poor growth under culturing conditions, without change of pH.

**Table 2.** Protologue for *Allorhizobium ampelinum* sp. nov.

|  |  |
| --- | --- |
| **Species name** | *Allorhizobium ampelinum* |
| **Genus name** | *Allorhizobium* |
| **Specific epithet** | *ampelinum* |
| **Species status** | sp. nov. |
| **Species etymology** | am.pe.li'num. Gr. n. ampelos grapevine; Gr. adj. ampelinos and<br>N.L. neut. adj. ampelinum of the vine |
| **Designation of the type strain** | S4 |
| **Strain collection numbers** | DSM 112012<sup>T</sup>, ATCC BAA-846<sup>T</sup> |
| **16S rRNA gene accession number** | U28505.1 |
| **Genome accession number** | GCF_000016285.1 |
| **Genome status** | Complete |
| **Genome size** | 6,320,946 |
| **GC mol %** | 57.47 |
| **Country of origin** | Hungary |
| **Region of origin** | Orgovány, Bács-Kiskun county |
| **Date of isolation** | 1981 |
| **Source of isolation** | Aerial gall on 2-year-old woody grapevine (*Vitis vinifera* cv. 'Izsaki<br>Sarfeher') |
| **Sampling date** | 1981 |
| **Number of strains in study** | 8 |
| **Source of isolation of non-type strains** | Grapevine |
| **Growth medium, incubation conditions used for standard cultivation** | Yeast mannitol agar (YMA) at 28 °C |
| **Conditions of preservation** | -80°C |
| **Gram stain** | Negative |
| **Cell shape** | Rod |
| **Colony morphology** | Colonies on YMA are white to cream coloured, circular, convex and glistening |
| **Positive tests with BIOLOG** | pH 6, D-Mannose, D-Galactose, 1% Sodium Lactate, Pectin, Rifamycin SV, Tetrazolium Blue, Potassium Tellurite |
| **Negative tests with BIOLOG** | pH5, N-Acetyl Neuraminic Acid, 4% NaCl, 8% NaCl, 3-Methyl Glucose, Inosine, Fusidic Acid, Troleandomycin, D-Serine, Minocycline, Guanidine HCl, Niaproof 4, p-Hydroxy-Phenylacetic Acid, Lithium Chloride, γ-Amino-Butyric Acid, α-Hydroxy-Butyric Acid, α-Keto-Butyric Acid, Propionic Acid, Sodium Butyrate |
| **Positive tests with API** | URE, ESC, PNG, GLU (assimilation), ARA, MNE, MAN, MLT, OX |
| **Negative tests with API** | NO<sub>3</sub>, TRP, GLU (fermentation), ADH, GEL, CAP, ADI, PAC |
| **Variable tests with API** | NAG, MAL, GNT, CIT |
| **Commercial kits used** | BIOLOG GEN3, API 20NE |
| **Oxidase<sup>a</sup>** | Positive |
| **Positive tests<sup>a</sup>** | Growth at 35°C, growth in nutrient broth supplemented with 2% NaCl, citrate utilization, production of acid from sucrose, production of alkali from tartrate |
| **Negative tests<sup>a</sup>** | Production of 3-ketolactose from lactose, acid-clearing on PDA with CaCO<sub>3</sub>, production of reddish-brown pellicle at the surface of ferric ammonium citrate broth, motility at pH 7.0, acid from d-(+)-melezitose, acid production from 4-hydroxyphenylacetate |
| **Known pathogenicity** | Plant pathogenic |

<sup>a</sup>These tests were performed for strains S4<sup>T</sup>, KFB 243, KFB 250, KFB 254 and KFB 264 by (42), except for a test of acid production in a medium containing 4-hydroxyphenylacetate conducted in this study. For strains that were not included in our former study, test of production of alkali from tartrate was conducted in the present work.

Although *All. vitis sensu stricto* strains carry genes predicted to be involved in a degradation process of gentisic acid, this biochemical property could not be demonstrated in this study. Gentisic acid degradation genes could have lost their function or not be induced under our test conditions. Alternatively, the predicted function might be incorrect and the target substrate of these enzymes may be an unidentified compound more or less closely related to gentisic acid.

We also tested the ability of *All. vitis* species complex strains to metabolize L-tartaric acid and produce alkali from this compound. In the present study, we included only strains that were not tested in our former work (42). Taken together, all tested *All. vitis* species complex strains (Table 1) were able to produce alkali from tartrate. Interestingly, strains IPV-BO 6186 and IPV-BO 7105, for which we could not identify TAR gene clusters, were also positive for this test.

As a broader way to characterize and phenotypically distinguish strains, we used MALDI-TOF mass-spectrometry (MS) of pure bacterial cultures. MALDI-TOF MS revealed diversity among the tested strains, while allowing to discriminate genomic species (Fig. S5).

### Relationship of the genus *Allorhizobium* and related *Rhizobiaceae* genera

As indicated by the core-genome phylogeny, the genus *Allorhizobium* is clearly separated from the other representatives of the family *Rhizobiaceae*, including the “*R. aggregatum* complex”, which, with the genus *Ciceribacter*, formed a well-delineated sister clade to *Allorhizobium* clade (Figs. 1, S1 and S2). The genome-based comparisons showed a clear divergence between these two clades. In particular, members of the genus *Allorhizobium* shared >74.9% AAI among each other, and 70.79-72.63% AAI with members of the “*R. aggregatum* complex”/*Ciceribacter* clade (Table S7). On the other hand, representatives of the genera *Shinella, Ensifer* and *Pararhizobium* showed 71.46-75.85% AAI similarity between genera. Similarly, representatives of genera *Neorhizobium* and *Pseudorhizobium* showed 72.24-76.18 % AAI similarity between genera. In other words, AAI values suggested that the existing genera *Ensifer, Pararhizobium* and *Shinella*, or *Neorhizobium* and *Pseudorhizobium* were more closely related than the genus *Allorhizobium* and the “*R. aggregatum* complex”/*Ciceribacter* clade. gANI and POCP values similarly supported the divergence of the members of *Allorhizobium* genus and the “*R. aggregatum* complex”/*Ciceribacter* clade (Table S7). Members of the genus *Allorhizobium* exhibited gANI and POCP values ranging 73.55-76.86 and 55.27-66.17, respectively, when compared with members of the “*R. aggregatum* complex”/*Ciceribacter* clade, values that were similar to these seen between representatives of the genera *Agrobacterium* and *Neorhizobium* (gANI 74.66-77.45; POCP 59.96-65.58).

## Discussion

### *Allorhizobium vitis* is not a single species

Genomic analyses allowed us to unravel the substantial taxonomic diversity within *All. vitis*. In particular, whole-genome sequence comparisons and phylogenomic analyses clearly showed that *Allorhizobium vitis* is not a single species, but represents a species complex composed of several genomic species. Similarly, *Agrobacterium* biovar 1 (i.e. *A. tumefaciens*) was initially considered a single species, but was later designated as a species complex comprising closely related, but distinct genomic species. Several studies applying DDH initially demonstrated this species diversity within *Agrobacterium* biovar 1 (43-45), which was later supported by results obtained with AFLP (46, 47), housekeeping gene analysis (10, 11, 14) and whole-genome sequence analysis (30). Although Ophel and Kerr (9) also performed DDH for several *All. vitis* strains, diversity within this species remained unknown because these authors only studied strains that belonged to *All. vitis sensu stricto* as defined here.

Our previous study based on the analysis of several housekeeping gene sequences suggested the existence of several phylogenetic groups within *All. vitis* species complex (28). The present study focused on two phylogenetic groups defined in our previous study: the first comprises the type strain of *All. vitis* (strain K309^T^) (9, 48), whereas the second includes the well-characterized and completely sequenced strain S4^T^ (18). Consequently, we amended the description of *All. vitis*, which now refers to the limited group within *All. Vitis* species complex strains (*All. vitis sensu stricto*) and proposed a description of a novel species, *All. ampelinum* sp. nov. (see formal description below).

As indicated by the genome analysis of a larger set of strains available from the NCBI GenBank database, the taxonomic diversity of *All. vitis* species complex is not limited to *All. vitis sensu stricto* and *All. ampelinum* sp. nov. However, the description of sub-clades C, D and E (Fig. S2) as separate species was considered outside the scope of this study, because the sequencing of these strains was not conducted by our group and their draft genome sequences are yet to be described in scientific publication(s). In addition, it is not clear whether sub-clades C and D represent a single or separate species. Further comprehensive genomic analysis of diverse members of these clades is required to elucidate relationships between them.

### Specific functions and ecologies suggested by clade-specific gene cluster analysis

The convergence of functions encoded by the *All. vitis* species complex-specific genes suggests an ancient adaptation to different kind of stresses, including exposure to aromatic compounds, competition with other rhizospheric bacteria and pH change. The occurrence of multiple signal perception systems in the *All. vitis* species complex-specific gene set indicates that adaptation to a changing environment is to be a key feature of their ecology.

We also searched genomes of *All. vitis* species complex strains for genes and gene clusters that were previously reported as important for the ecology of this bacterium. In this regard, polygalacturonase production, a trait associated with grapevine root necrosis (34, 49, 50), and tartrate degradation (37) were proposed to contribute to the specialization of *All. vitis* to its grapevine host. In addition, polygalacturonase activity might be involved in the process of the invasion of the host plant, as postulated previously for other rhizobia (51). Although all *All. vitis* species complex strains carried the *pehA* gene encoding a polygalacturonase enzyme, this gene was not restricted to this bacterial group, as it was also present in all other *Allorhizobium* spp. strains included in our analysis, except for *All. undicola*.

All *All. vitis* species complex strains included in this study, except for strains IPV-BO 6186 and IPV-BO 7105, carried TAR regions. However, all of them were able to metabolize tartrate and produce alkali from this compound. Therefore, we speculate that strains IPV-BO 6186 and IPV-BO 7105 must carry another type of TAR system, distinct from those described so far in other *All. vitis* strains. Furthermore, some diversity between TAR regions and variable distribution patterns of different TAR regions among strains were observed, in line with previously reported data (39). The existence of non-tartrate-utilizing strains was also documented in the literature (39). Considering the fact that tartrate utilization in *All. vitis* has only been observed as plasmid-borne (40, 41, 52), this suggests that tartrate utilization is an accessory trait that can be readily gained via the acquisition of a plasmid encoding this trait and selected for in tartrate-abundant environments. Because grapevine is rich in tartrate (53), utilization of this substrate may enhance the competitiveness of *All. vitis* species complex strains in colonizing this plant species (37).

We observed that an important fraction of the species-specific genes for *All. vitis sensu stricto* and *All. ampelinum* occurred on chromids and plasmids, suggesting that these replicons may be an important part of these species’ adaptive core-genome, as previously observed in the *A. tumefaciens* species complex (30). Ecological differentiation of the two main species of the *All. vitis* species complex seems to rely on consumption of different nutrient sources, including polyamines and nickel ion (potentially as a key cofactor of ecologically important enzymes) for *All. ampelinum*, and phenolic compounds for *All. vitis sensu stricto*.

Even though *All. vitis sensu stricto* strains carried a putative gene cluster which predicted function was the degradation of gentisic acid, we could not experimentally demonstrate this trait. Gentisic acid was detected in grapevine leaves (54) and is likely present in other parts of this plant. This compound was reported as a plant defense signal that can accumulate in some vegetable plants responding to compatible viral pathogens (55, 56). In addition, a sub-clade within *All. vitis sensu stricto* composed of strains K309^T^ and KFB 253 carried a complete pathway for degradation of salicylic acid through gentisic acid. Salicylic acid is recognized as an important molecule for plant defense against certain pathogens (57). The role of salicylic and gentisic acid in grapevine defense mechanism against pathogenic bacteria has not been studied in detail, and further investigations are required to understand their effect against tumorigenic agrobacteria. Furthermore, we predicted, and demonstrated that all studied *All. vitis sensu stricto* strains have the specific ability to degrade 4-hydroxyphenylacetate, an activity that may contribute to the detoxication of aromatic compounds and thus to the survival of this bacterium in soil, notably in competition against bacteria lacking this pathway.

Similarly, gene clusters putatively involved in polyamine metabolism or nickel assimilation might confer to *All. ampelinum* the ability to persist in harsh environments. In this respect, nickel import has been shown to be essential for hydrogenase function in *E. coli* (58). Hydrogenase function has in turn been proposed as a potential mechanism for detoxication of phenolic compounds in *A. vitis* (59) and may thus have an important role in survival in the rhizosphere.

### Delineation of the genus *Allorhizobium*

The genus *Allorhizobium* was clearly differentiated from other *Rhizobiaceae* genera based on core- and pan-genome-based phylogenies, in line with previous studies employing genome-wide phylogeny (15, 16). We included diverse *All. vitis* species complex strains into our analysis, confirming that these bacteria, principally recognized as grapevine crown gall causative agents, belong to the genus *Allorhizobium*.

On the other hand, the taxonomic status of the “*R. aggregatum* complex”/*Ciceribacter* clade is still unresolved. Although MLSA suggested that “*R. aggregatum* complex” is a sister clade of the genus *Agrobacterium*, the more thorough phylogenetic analyses performed in this study rather showed that the “*R. aggregatum* complex” grouped with *Ciceribacter* spp., in a clade that is more closely related to the genus *Allorhizobium*. Presently, there is no widely accepted criteria and scientific consensus regarding the delineation of new bacterial genera (27). In this study, existing *Rhizobiaceae* genera were compared using several delineation methods proposed in the literature, such as AAI (60, 61), POCP (62) and gANI/AF (63), which we complemented with genome-based phylogenies. Taken together, our genome-based analysis suggested that *Allorhizobium* represents a genus clearly separated from other *Rhizobiaceae* genera, including closely related “*R. aggregatum* complex”/*Ciceribacter* clade. A separate and more focused analysis is however required to explore the taxonomic diversity and structure of the “*R. aggregatum* complex”/*Ciceribacter* clade.

## Conclusions

Whole-genome sequence comparisons and phylogenomic analyses placed *All. vitis* strains within the genus *Allorhizobium*, which was clearly differentiated from other *Rhizobiaceae* genera, including the closely related “*R. aggregatum* complex”/*Ciceribacter* clade. We revealed an extensive and structured genomic diversity within *All. vitis*, which in fact represents a species complex composed of several genomic species. Consequently, we emended the description of *All. vitis*, now encompassing a restricted group of strains within the *All. vitis* species complex (i.e. *All. vitis sensu stricto*) and proposed a description of a novel species, *All. ampelinum* sp. nov. Further analyses including pan-genome reconstruction and phylogeny-driven comparative genomics revealed loci of genomic differentiation between these two species. Functional analysis of these species-specific loci suggested that these species are ecologically differentiated as they can consume specific nutrient sources (*All. ampelinum*), or degrade specific toxic compounds (*All. vitis sensu stricto*). We identified another two potential genomic species within the *All. vitis* species complex, for which the limited diversity of available isolates prevented further characterization. We also described how accessory genomic regions associated with the colonization of grapevine host plant are distributed across species, and how they combine to form diverse genotypes. However, given the complete bias in sampling of *All. vitis* strains – all grapevine pathogens – the ecological significance of this genetic diversity remains unclear. We encourage future studies to integrate genomic data from new genomically diverse isolates, to further unravel the ecological basis of *All. vitis* species complex diversification.

### Emended description of *Allorhizobium vitis* (Ophel and Kerr 1990) Mousavi et al. 2016 emend. Hördt et al. 2020

The description of *Agrobacterium vitis* is provided by Ophel and Kerr (9). Young et al. (13) proposed the transfer of *A. vitis* to the genus *Rhizobium*, but it was neither widely accepted by the scientific community nor supported by further studies (14, 64, 65). Mousavi et al. (11) reclassified this species to the genus *Allorhizobium*, which was included into the Validation list no. 172 of the IJSEM (66). Hördt et al. (16) emended a description of *All. vitis* by including genome sequence data for its type strain, which was published in the List of changes in taxonomic opinion no. 32 (67).

As shown in this study, *All. vitis sensu stricto* includes a limited group of strains that can be differentiated from other *All. vitis* genomic species and other *Allorhizobium* species based on OGRIs, such as ANI, as well as by core-genome phylogeny. Moreover, *All. vitis sensu stricto* can be differentiated from other species of *All. vitis* species complex by analysis of sequences of housekeeping genes *dnaK, gyrB* and *recA* (28). Finally, this study demonstrated that strains belonging to this species can be distinguished from *All. ampelinum* by MALDI-TOF MS analysis. Unlike any *All. ampelinum*, all tested *All. vitis sensu stricto* strains are able to produce acid in a medium containing 4-hydroxyphenylacetate. However, this apparently species-specific trait is borne by a plasmid, and could possibly be transmitted to closely related species.

The whole-genome sequence of type strain K309 is available in GenBank under the accessions LMVL00000000.2 and GCA_001541345.2 for the Nucleotide and Assembly databases, respectively (48). The genomic G+C content of the type strain is 57.55%. Its approximate genome size is 5.75 Mbp.

Basonym: *Agrobacterium vitis* Ophel and Kerr 1990.

The type strain, K309^T^ (= <u>NCPPB 3554</u>^T^ =HAMBI 1817^T^ = <u>ATCC 49767</u>^T^ = CIP 105853^T^ = <u>ICMP 10752</u>^T^ = <u>IFO 15140</u>^T^ = <u>JCM 21033</u>^T^ = <u>LMG 8750</u>^T^ = <u>NBRC 15140</u>^T^), was isolated from grapevine in South Australia in 1977.

### Description of *Allorhizobium ampelinum* sp. nov

The description and properties of the new species are given in the protologue (Table 2).

*All. ampelinum* (am.pe.li’num. Gr. n. ampelos grapevine; Gr. adj. ampelinos and N.L. neut. adj. *ampelinum* of the vine).

*All. ampelinum* strains were formerly classified in the species *All. vitis*. However, our genomic data showed that they can be distinguished from *All. vitis sensu stricto* and other *All. vitis* genomic species based on OGRIs (e.g. ANI and dDDH) and core-genome phylogeny, as well as by analysis of sequences of housekeeping genes (28). Furthermore, *All. ampelinum* can be differentiated from *All. vitis sensu stricto* by MALDI-TOF MS analysis.

The type strain, S4^T^ (= DSM 112012^T^ = <u>ATCC BAA-846</u>^T^) was isolated from grapevine tumor in Hungary in 1981.

## Methods

### *Allorhizobium vitis* strains

*All. vitis* strains used in this study were isolated from crown gall tumors on grapevine originating from different geographical areas (Table 1). These strains were predominantly representatives of the two main phylogenetic groups (C and D) delineated in our previous study (28).

### DNA extraction

For whole genome sequencing, genomic DNA was extracted from bacterial strains grown on King’s medium B (King et al. 1954) at 28°C for 24 h using NucleoSpin Microbial DNA kit (Macherey-Nagel, Germany). The quality of the genomic DNA was assessed by electrophoresis in 0.8% agarose gel.

### Genome sequencing

Draft whole-genome sequences were obtained for 11 *All. vitis* strains (Table 1). DNA libraries were obtained with Nextera XT DNA Library Prep Kit (Illumina, USA). Paired-end sequencing (2 × 300 bp) was performed on an Illumina MiSeq platform generating 487,883-2,309,377 paired reads per genome. Trimming and quality filtering of raw reads were conducted using Trimmomatic (Galaxy Version 0.36.5) (68) implemented on the Galaxy Web server (69). The read quality was assessed with FastQC (Galaxy Version 0.72+galaxy1) (http://www.bioinformatics.babraham.ac.uk/projects/fastqc/). In order to achieve higher coverage for strains Av2, IPV-BO 1861-5, KFB239 and KFB 264, additional paired-end sequencing (2 × 150 bp) was performed using an Illumina NextSeq 500 platform generating 1,037,619-1,443,575 paired reads. Demultiplexing and adapter clipping was done using the bcl2fastq(2) conversion software (Illumina, USA).

### Genome assembly and annotation

*De novo* genome assemblies were performed using the SPAdes genome assembler (Galaxy Version 3.12.0+galaxy1) (70). For genomes sequenced on the MiSeq and NextSeq platforms, both sets of reads were used for assembly. The genome sequences were deposited to DDBJ/ENA/GenBank under the Whole Genome Shotgun projects accession numbers listed in Table 1, under BioProject ID PRJNA557463.

The genome sequences were annotated using Prokka (Galaxy Version 1.13) (71) and NCBI Prokaryotic Genomes Annotation Pipeline (PGAP) (72). Prokka Version 1.14.6 was used to annotate genomes as a part of the Pantagruel pipeline (task 0; see below and Supplementary Methods). Functional annotation of proteins encoded by each gene family clustered by Pantagruel was conducted by the InterProScan software package Version 5.42-78.0 (Jones et al., 2014) as implemented in the Pantagruel pipeline (Task 4). Additionally, annotation of particular sequences of interest and metabolic pathway prediction were performed using BlastKOALA and GhostKOALA (last accessed on December, 2020) (73). Protein sequences analyzed were subjected to Pfam domain searches (database release 32.0, September 2018, 17929 entries) (74). Metabolic pathway prediction was performed using KEGG (75) and MetaCyc (76) databases (last accessed on December, 2020).

The NCBI BLASTN and BLASTP (https://blast.ncbi.nlm.nih.gov/Blast.cgi), as well as BLAST search tool of KEGG database (last accessed on December, 2020) (75), were used for ad-hoc sequence comparisons at the nucleotide and amino acid levels, respectively.

### Core- and pan-genome phylogenomic analyses

For phylogenomic analysis, whole genome sequences of 69 *Rhizobiaceae* strains were used, including 14 strains of *All. vitis* (Table 1) and 55 reference *Rhizobiaceae* strains (Table S1a). Additionally, in order to further explore the phylogenetic diversity of *All. vitis*, another core-genome phylogeny was inferred from an extended dataset that also included 34 *All. vitis* genomes available from GenBank but not yet published in peer-review journals by sequence depositors (Table S1b). To build phylogenies based on the core-genome (supermatrix of concatenated non-recombining core gene alignments) and on the pan-genome (homologous gene cluster presence/absence matrix), we used the GET_HOMOLOGUES Version 10032020 (33) and GET_PHYLOMARKERS Version 2.2.8.1_16Jul2019 (77) software packages. Details of the bioinformatic pipeline and used options are described in the Supplementary Methods.

### Overall genome relatedness indices

To differentiate between the strains, different OGRIs were computed. For species delimitation, we relied on the values of average nucleotide identity (ANI) (29, 78) and digital DNA-DNA hybridization (dDDH) (79) among strain genomes. Because different implementations of the ANI metric are known to give slightly different results (80), ANI was calculated using several programs: PyANI Version 0.2.9 (for metrics ANIb and ANIm) (81) (https://github.com/widdowquinn/pyani), OrthoANIu Version 1.2 (82) and FastANI Version 1.2 (83) tools. dDDH values were calculated using the Genome-to-Genome Distance Calculator (GGDC) Version 2.1 (79).

For genus delimitation, we relied on average amino acid identity (AAI) (22, 60, 78), genome-wide average nucleotide identity (gANI) and alignment fraction (AF) (84), and the percentage of conserved proteins (POCP) (62). AAI values were calculated with CompareM Version 0.0.23 (https://github.com/dparks1134/CompareM). gANI and AF values were obtained by the ANIcalculator Version 1.0 (84). POCP values were calculated using GET_HOMOLOGUES software package (33). Details of the used software and options are given in the Supplementary Methods.

### Genome gene content analyses and identification of clade-specific genes

To explore the distribution of genome gene contents, we conducted further pan-genome analyses on more focused datasets, using two different bioinformatics pipelines, from which we present a consensus. Firstly, a pan-genome database was constructed using the Pantagruel pipeline Version 00aaac71f85a2afa164949b86fbc5b1613556f36 under the default settings as described previously (31, 32) and in Supplementary Methods. Because of computationally intensive tasks undertaken in this pipeline, the dataset was limited to the *Allorhizobium* genus and its sister clade “*Rhizobium aggregatum* complex”/*Ciceribacter* (28 strains).

Secondly, we analyzed a more focused dataset comprised of the 14 *All. vitis* species complex strains (Table 1) and four *Allorhizobium* spp. (*All. oryziradicis* N19^T^, *All. taibaishanense* 14971^T^, *All. terrae* CC-HIH110^T^ and *All. undicola* ORS 992^T^; Table S1a), using the GET_HOMOLOGUES software package (33). Pan-genome gene clusters were classified into core, soft core, cloud and shell compartments (85) and species-specific gene families were identified from the pan-genome matrix. For details on the used scripts and options, see Supplementary Methods.

### Biochemical tests

*All. vitis* strains were phenotypically characterized using API and Biolog tests. The API 20NE kit was used according to manufacturer’s instructions (bioMérieux). Utilization of sole carbon sources was tested with Biolog GEN III microplates using protocol A, according to the instructions of the manufacturer (Biolog, Inc., Hayward, CA, USA).

The metabolism of 4-hydroxyphenylacetic acid (p-hydroxyphenylacetic acid; Acros Organics, Product code: 121710250) and gentisic acid (2,5-dihydroxybenzoic acid; Merck, Product Number: 841745) was performed in AT minimal medium (86, 87) supplemented with yeast extract (0.1 g/L), bromthymol blue (2.5 ml/L of 1 % [w/v] solution made in 50% ethanol), and the tested compound (1 g/L). Hydroxyphenylacetic and gentisic acid were added as filter-sterilized 1% aqueous solutions. Bacterial growth and color change of the medium were monitored during one week of incubation at 28°C and constant shaking (200 rpm/min). Metabolism of L(+)-tartaric acid, involving production of alkali from this compound, was tested as described before (5).

### MALDI-TOF Mass Spectrometry analysis

Sample preparation for MALDI-TOF mass spectrometry (MS) was carried out according to the Protocol 3 described by Schumann and Maier (88). Instrument settings for the measurements were as described previously by Tóth et al. (89). The dendrogram was created using the MALDI Biotyper Compass Explorer software (Bruker, Version 4.1.90).

## Abbreviations

AAI: Average amino acid identity
AF: Alignment fraction
ANI: Average nucleotide identity
DDH: DNA-DNA hybridization
dDDH: Digital DDH
gANI: Genome-wide ANI
MS: Mass-spectrometry
OGRI: Overall genome relatedness index
MLSA: Multi-locus sequence analysis
PGAP: Prokaryotic genomes annotation pipeline
POCP: Percentage of conserved proteins
TAR: Tartrate utilization
Ti: Tumor-inducing

## Acknowledgments

We would like to thank Cathrin Spröer and Boyke Bunk for conducting the Illumina sequencing. We are grateful to Anja Frühling and Ulrike Steiner for support in phenotypic tests.

## Authors’ contributions

NK and FL conceived and designed the study, and analyzed data. FL conceived and implemented the bioinformatic analysis pipeline Pantagruel. SV performed phenotypic characterization of the strains (API, Biolog and MALDI-TOF). EB provided bacterial strains. JP, JO and KS were involved in interpreting data. NK and FL wrote the manuscript. All authors read, discussed, edited and approved the final manuscript.

## Funding

This research was supported by the Georg Forster Fellowship for postdoctoral research from the Alexander von Humboldt-Foundation, Bonn, Germany. The work of FL was supported by by a Medical Research Council (MRC) grant (<u>MR/N010760/1</u>) and the Wellcome Trust Grant [206194]. Bioinformatic analyses were supported by the BMBF-funded de.NBI Cloud within the German Network for Bioinformatics Infrastructure (de.NBI) (031A537B, 031A533A, 031A538A, 031A533B, 031A535A, 031A537C, 031A534A, 031A532B).

## Availability of data and materials

The genome sequences generated in this study were deposited to DDBJ/ENA/GenBank under the Whole Genome Shotgun projects accession numbers listed in Table 1, under BioProject ID PRJNA557463. The versions described in this paper are first versions.

All other relevant data (including output of analyses) referring to this project have been deposited on Figshare under the project accession 20894, available at figshare (https://figshare.com/), with individual items accessible at DOIs: 10.6084/m9.figshare.17105267, 10.6084/m9.figshare.17125571, 10.6084/m9.figshare.16850071, 10.6084/m9.figshare.16849165, 10.6084/m9.figshare.13440218 and 10.6084/m9.figshare.17125568.

(DOIs to be published upon manuscript acceptance; private links for peer-review: https://figshare.com/s/b6c7ccc50f0179a08ce6, https://figshare.com/s/6aaae45f5eb5a81baeb5, https://figshare.com/s/c88e7412cf58bc9b1b54, https://figshare.com/s/77bb93622b1bf788bada, https://figshare.com/s/f17bb1c00a960714e097, https://figshare.com/s/eecc49ea166bbe061554).

## Declarations

### Ethics approval and consent to participate

Not applicable.

### Consent for publication

Not applicable.

### Competing interests

The authors declare no competing interests.

## Supplementary Information

**Additional file 1: Fig. S1**. Maximum-likelihood core-genome phylogeny of 69 strains belonging to the genus *Allorhizobium* and other *Rhizobiaceae* members (uncollapsed). The tree was estimated with IQ-TREE from the concatenated alignment of 344 top-ranked genes selected using GET_PHYLOMARKERS software. The numbers on the nodes indicate the approximate Bayesian posterior probabilities support values (first value) and ultra-fast bootstrap values (second value), as implemented in IQ-TREE. The tree was rooted using the *Mesorhizobium* spp. sequences as the outgroup. The scale bar represents the number of expected substitutions per site under the best-fitting GTR+F+ASC+R6 model. The same tree, but with collapsed clades, is presented in Figure 1.

**Additional file 2: Fig. S2**. Maximum-likelihood core-genome phylogeny of 103 strains belonging to the genus *Allorhizobium* (including 34 additional strains of *All. vitis* species complex strains whose sequences are available in GenBank but not associated to a published study) and other *Rhizobiaceae* members. The tree was estimated with IQ-TREE from the concatenated alignment of 302 top-ranked genes selected using GET_PHYLOMARKERS software. The numbers on the nodes indicate the approximate Bayesian posterior probabilities support values (first value) and ultra-fast bootstrap values (second value), as implemented in IQ-TREE. The tree was rooted using the *Mesorhizobium* spp. sequences as the outgroup. The scale bar represents the number of expected substitutions per site under the best-fitting GTR+F+ASC+R7 model. The matrix in the top-right corner represents the distribution of ANIb values for genomic sequences of the clade corresponding to the *All. vitis* species complex, relative to the typical species delimitation threshold of 95%.

**Additional file 3: Fig. S3**. Maximum-likelihood pan-genome phylogeny of 69 strains belonging to the genus *Allorhizobium* and other *Rhizobiaceae* members (uncollapsed). The tree was estimated with IQ-TREE from the consensus (COGtriangles and OMCL clusters) pan-genome matrix containing 33,396 clusters obtained using GET_HOMOLOGUES software. The numbers on the nodes indicate the approximate Bayesian posterior probabilities support values (first value) and ultra-fast bootstrap values (second value), as implemented in IQ-TREE. The tree was rooted using the *Mesorhizobium* spp. sequences as the outgroup. The scale bar represents the number of expected substitutions per site under the best-fitting GTR2+FO+R5 model. The same tree, but with collapsed clades, is presented in Figure 2.

**Additional file 4: Fig. S4**. Heatmap representation of the average nucleotide identity (ANIb) for TAR regions of *All. vitis* species complex strains. PyANI program Version 0.2.9 (https://github.com/widdowquinn/pyani) was used to calculate ANIb values and generate the clustered heatmap.

**Additional file 5: Fig. S5**. Score-oriented dendrogram showing the similarity of the MALDI-TOF mass spectra of 14 *All. vitis* species complex strains studied. The dendrogram was created using the MALDI Biotyper Compass Explorer software (Bruker, Version 4.1.90).

**Additional file 6: Table S1**. List of additional strains and GenBank/EMBL/DDBJ accession numbers for their nucleotide sequences used in this study. **a)** List of 55 reference *Rhizobiaceae* strains and GenBank/EMBL/DDBJ accession numbers for their nucleotide sequences used in this study; **b)** List of additional 34 *All. vitis* species complex strains and GenBank/EMBL/DDBJ accession numbers for their nucleotide sequences used in this study. Although available in the public nucleotide sequence databases, these genome sequences have not yet been presented in peer-reviewed study by sequence depositors.

**Additional file 7: Table S2**. Pairwise OGRI comparisons amongst 14 *Allorhizobium vitis* species complex strain genomes towards species delimitation. ANI values were computed with various implementation of the ANI metric: **a)** ANIb, **b)** ANIm, **c)** orthoANI and **d)** fastANI; **e)** dDDH values were computed with the GGDC.

**Additional file 8: Table S3**. Pairwise ANIb comparisons amongst extended set of 48 *Allorhizobium vitis* species complex strain genomes towards species delimitation.

**Additional file 9: Table S4**. Clusters of contiguous clade-specific genes. Clusters were identified amongst sets of genes deemed specific of the focal clade based on detection by either Pantagruel or GET_HOMOLOGUES pipelines. **a)** Clusters of genes specific to *Allorhizobium vitis* species complex (present in all *All. vitis sensu stricto, All. ampelinum* and *Allorhizobium* sp. Av2, and in no other *Allorhizobium* spp.); **b)** Clusters of genes specific to *All. vitis sensu stricto* (present in all five tested strains and in none of *All. ampelinum*); **c)** Clusters of genes specific to *All. ampelinum* (present in all eight tested strains and in none of *All. vitis sensu stricto*).

**Additional file 10: Table S5**. Pairwise ANIb values between tartrate utilization (TAR) regions of *Allorhizobium vitis* species complex strains.

**Additional file 11: Table S6**. Tartrate utilization (TAR) region genotype and tartrate metabolism phenotype (production of alkali from L-tartaric acid) of *All. vitis* species complex strains.

**Additional file 12: Table S7**. Pairwise OGRI comparisons amongst 69 *Rhizobiaceae* strain genomes towards genus delimitation. **a)** Average amino acid identity (AAI); **b)** percentage of conserved proteins (POCP); **c)** genome-wide average nucleotide identity (gANI) and alignment fraction (AF), with AF values indicated in parentheses.

**Additional file 13: Supplementary Methods**. This file contains additional detailed methods description.

## Notes

https://figshare.com/projects/Allorhizobium_vitis/125029

### Competing Interest Statement

The authors have declared no competing interest.

### Summary of Updates

Title change; Abstract modified; Results section updated; Supplemental files updated;

