## Supplementary figures and images for "Genomic analysis provides novel insights into diversification and taxonomy of *Allorhizobium vitis* (i.e. *Agrobacterium vitis*)"

### Fig. S1

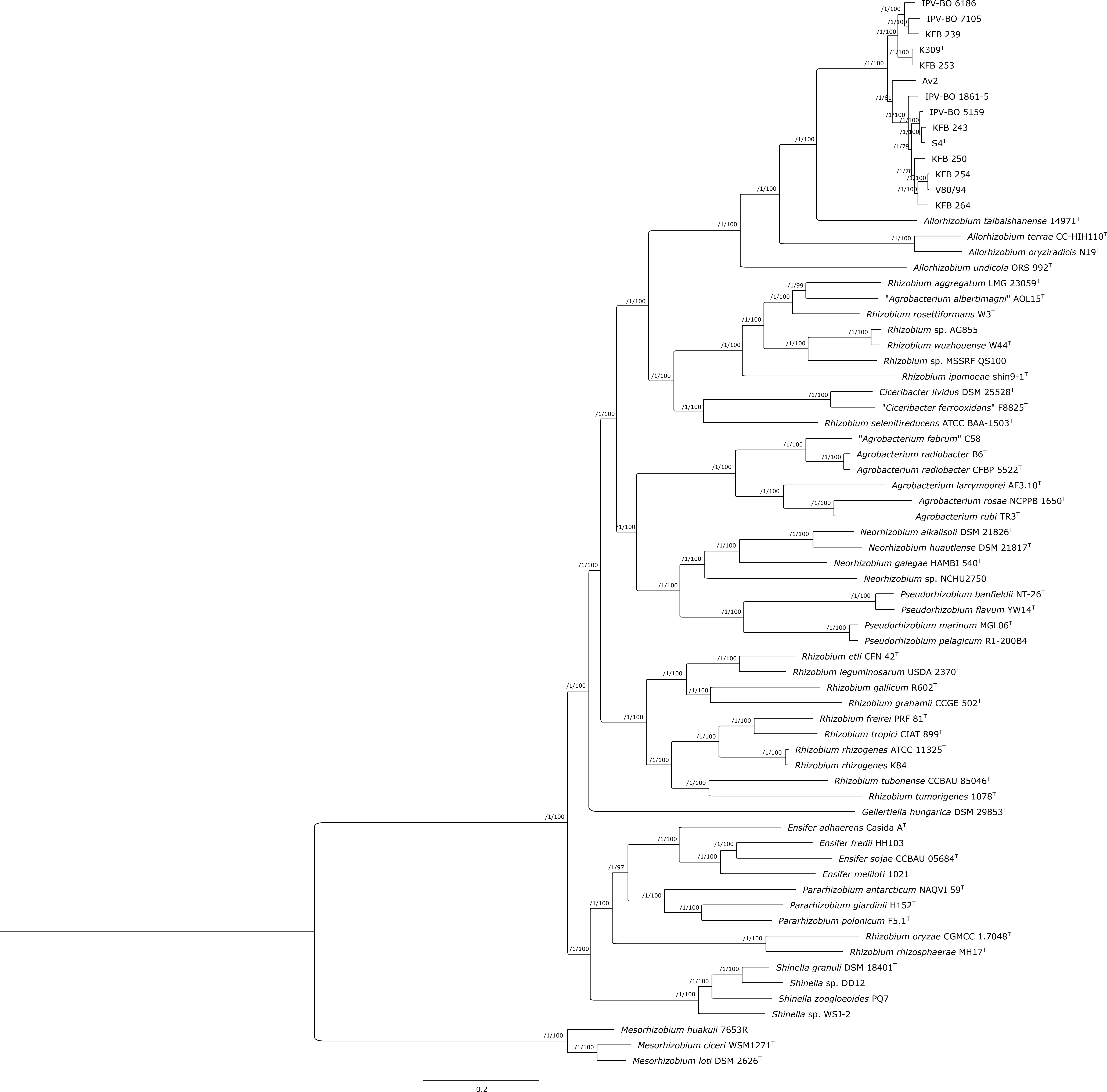

### Fig. S2

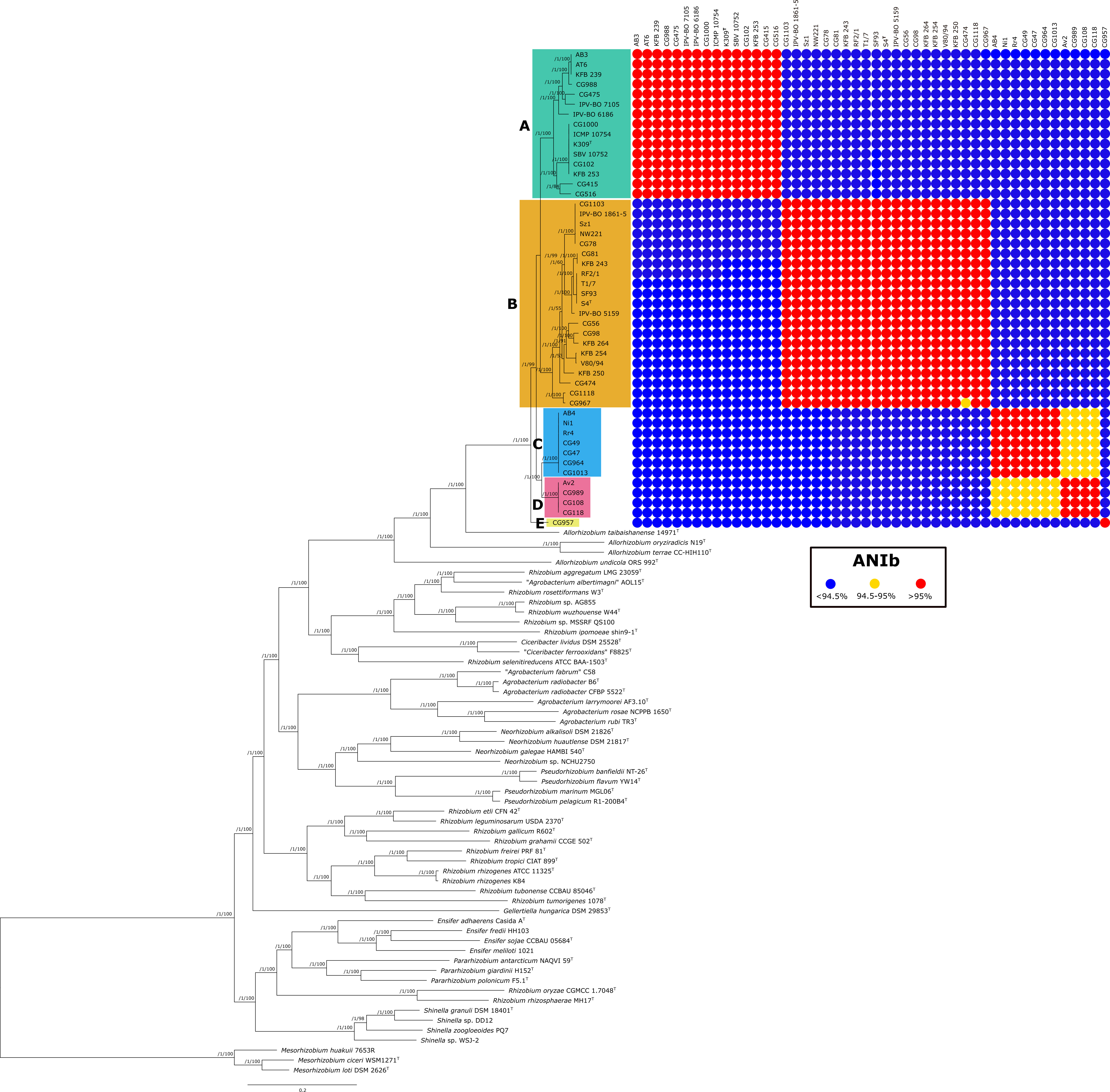

### Fig. S3

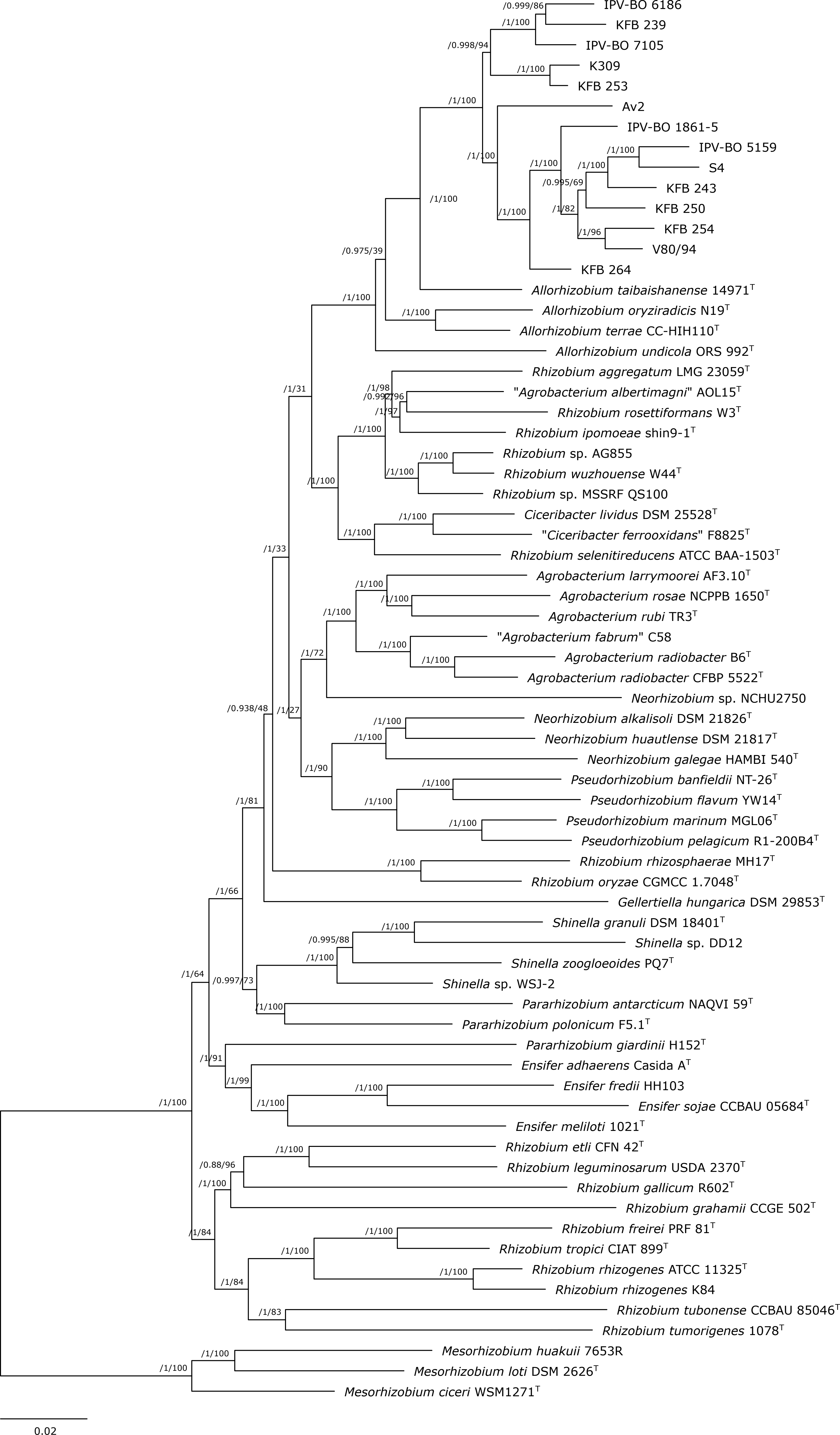

### Fig. S4

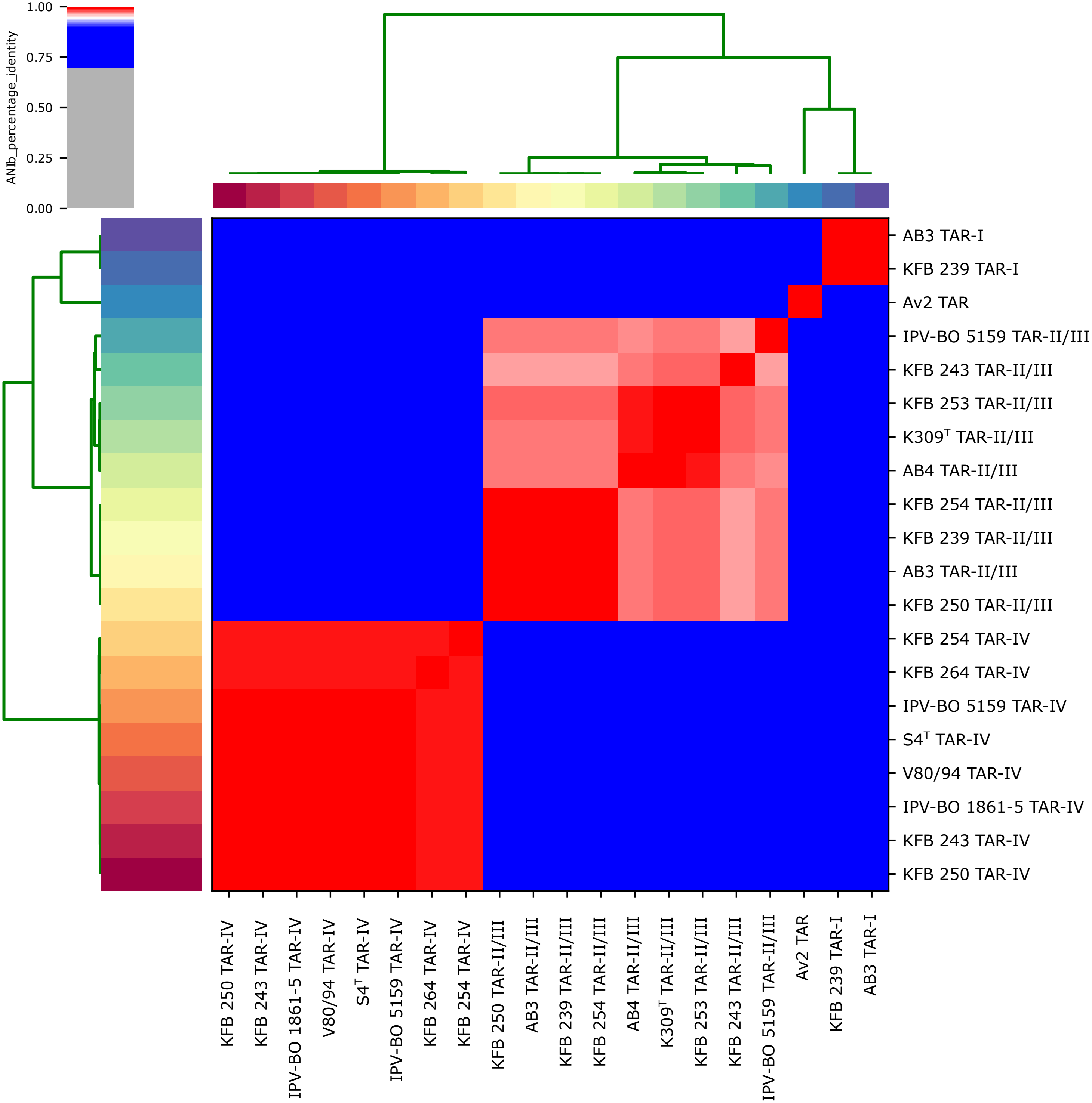

### Fig. S5

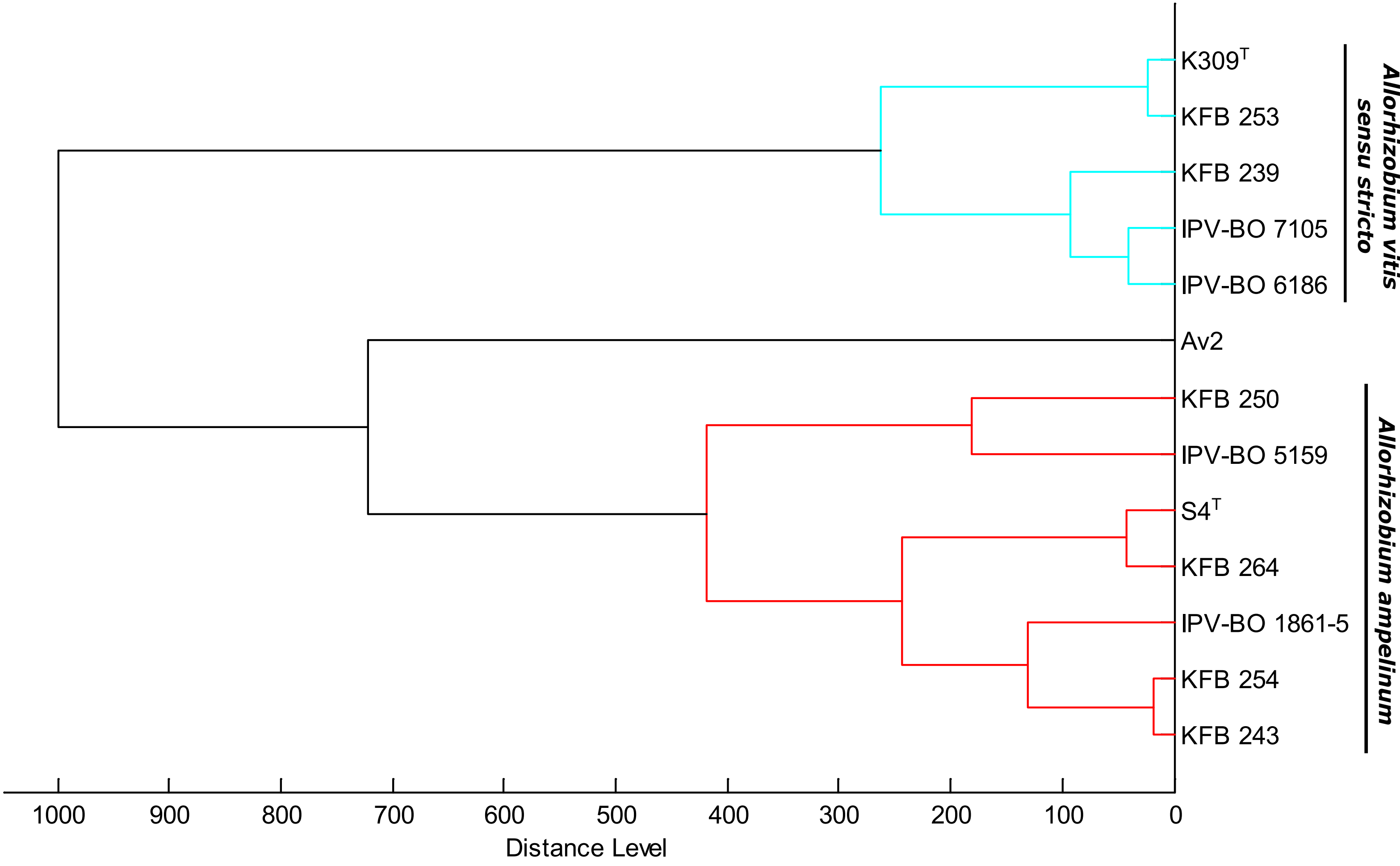
