## Supplementary Methods for "Genomic analysis provides novel insights into diversification and taxonomy of *Allorhizobium vitis* (i.e. *Agrobacterium vitis*)"

### Core- and pan-genome phylogenomic analyses

For phylogenomic analysis, whole genome sequences of 69 *Rhizobiaceae* strains were used, including 14 strains of *All. vitis* (Table 1) and 55 reference *Rhizobiaceae* strains (Table S1a). Additionally, in order to further explore the phylogenetic diversity of *All. vitis*, another core-genome phylogeny was inferred from an extended dataset that also included 34 *All. vitis* genomes available from GenBank but not yet published in peer-review journals by sequence depositors (Table S1b). For the core-genome phylogeny, GenBank files generated by Prokka were used as an input to GET_HOMOLOGUES software package version 10032020 (1). To compute homologous gene clusters, we ran the get_homologues.pl script, which implements the bidirectional best-hit (BDBH), Clusters of Orthologous Groups-triangles (COGtriangles), and OrthoMCL (Markov Clustering of orthologs, OMCL) algorithms. A stringent 90% coverage cut-off for BLASTP alignments (option “-C 90”) was imposed. A consensus core-genome was determined as the intersection of the clusters computed by the BDBH, COG-triangles and OMCL algorithms using the script compare_clusters.pl (option “-t *number of genomes*”). The resulting core-genome clusters were processed with the GET_PHYLOMARKERS software package version 2.2.8.1_16Jul2019 (2) with the options “-R 1 -t DNA”. This software was designed to select reliable phylogenetic markers and exclude sequences with the evidence of recombination, and those yielding anomalous or poorly resolved gene trees. The selected DNA sequences were aligned at the codon level and concatenated, producing a supermatrix alignment that were used for phylogenetic analysis. A maximum likelihood (ML) phylogeny was estimated under the best-fitting substitution model using IQ-TREE v1.6.10 (3) as implemented in the GET_PHYLOMARKERS package.

For the pan-genome phylogeny, pan-genome (presence-absence) matrices were computed from the consensus of homologous gene cluster sets determined by the COGtriangles and OMCL clustering algorithms (see above) using the compare_clusters.pl script (options “-t 0, -m”) from the GET_HOMOLOGUES software package. The ML pan-genome phylogeny was estimated from the pan-genome matrix (PGM) using the estimate_pangenome_phylogenies.sh script from the GET_PHYLOMARKERS software package. Twenty-five independent IQ-TREE searches fitting binary models were launched and the best fit was retained. Another PGM-based phylogeny based was inferred using parsimony with the script compare_clusters.pl (option –T) from the GET_HOMOLOGUES package.

### Overall genome relatedness indices

To differentiate the strains under study, different OGRIs were computed. For species delimitation, we relied on the values of average nucleotide identity (ANI) (4, 5) and digital DNA-DNA hybridization (dDDH) (6) among strain genomes. Because different implementations of the ANI metric are known to give slightly different results (7), ANI was calculated using several programs: PyANI Version 0.2.9, with scripts employing BLAST+ (ANIb) and MUMmer (ANIm) to align the input sequences) (8), OrthoANIu Version 1.2 (calculates orthologous ANI using USEARCH algorithm) (9) and FastANI Version 1.2 (estimates ANI using Mashmap as its MinHash-based alignment-free sequence mapping engine) (10) tools. dDDH values were calculated using the Genome-to-Genome Distance Calculator (GGDC 2.1) (6), using the recommended BLAST+ alignment and formula 2 (identities/HSP length).

For genus delimitation, we relied on average amino acid identity (AAI) (5, 11, 12) and genome-wide average nucleotide identity (gANI) and alignment fraction (AF) (13) and the percentage of conserved proteins (POCP) (14). AAI values were computed with CompareM Version 0.0.23 (<https://github.com/dparks1134/CompareM>) using default options. gANI and AF values were obtained with the ANIcalculator Version 1.0, where gANI computation involves comparison of orthologous protein-coding genes identified as bidirectional best hits (BBHs), at the nucleotide level, and AF is calculated as a fraction of the sum of the lengths of BBH genes divided by the sum of the lengths of all genes in a genome (13). For gANI/AF analysis, we used genes as annotated by Prodigal Version 2.6.3. POCP values were calculated using through GET_HOMOLOGUES software package (option “-P”) (1), based on the OMCL clustering of homologous genes (15).

### Genome gene content analyses and identification of clade-specific genes

To explore the distribution of genome gene contents, we conducted further pan-genome analyses on more focused datasets, using two different bioinformatics pipelines, from which we present a consensus. Firstly, a pan-genome database was constructed using the Pantagruel pipeline Version 00aaac71f85a2afa164949b86fbc5b1613556f36 under the default settings as described previously (16, 17) (<http://github.com/flass/pantagruel/>). Because of computationally intensive tasks undertaken in this pipeline, the dataset was limited to the *Allorhizobium* genus and its sister clade “*Rhizobium aggregatum* complex”/*Ciceribacter* (28 strains). Shortly, the Pantagruel pipeline was used to reconcile the topology of gene trees with the reference species tree (core-genome tree) for each gene family of the pan-genome that contained a least four sequences. Genes were classified into orthologous clusters based on their gain/loss scenarios, from which clade-specific gene sets were identified. Genes specific for a particular clade (species or genus) were identified from the gene count matrix output of Pantagruel task 08, using the R script get_clade_specific_genes.r available in Pantagruel. Over-representation of Gene Ontology functional annotation terms in clade-specific genes with respect to the clade’s core genomes was tested using the R script clade_specific_genes_GOterm_enrichment_test.r available in Pantagruel, using the enrichment test implemented in the topGO R package (18) with the ‘weight01’ algorithm and ‘Fisher’ statistics (tests with p-values lower than 0.1 were deemed significant).

Secondly, we analyzed a more focused dataset comprised of the 14 *All. vitis* species complex strains (Table 1) and four *Allorhizobium* spp. (*All. oryziradicis* N19^T^, *All. taibaishanense* 14971^T^, *All. terrae* CC-HIH110^T^ and *All. undicola* ORS 992^T^; Table S1a), using the GET_HOMOLOGUES software package (1). As described above, homologous gene clusters were computed using COGtriangles and OMCL algorithms, followed by computation of consensus pan-genome clusters and pan-genome matrix. Pan-genome gene clusters were classified into core, soft core, cloud and shell compartments (19) by the auxiliary script parse_pangenome_matrix.pl (option “-s”); accessory genes include both shell and cloud compartments. The script parse_pangenome_matrix.pl of GET_HOMOLOGUES software package was used to identify species-specific gene families (option “-g”) from the pan-genome matrix generated with the GET_HOMOLOGUES software package. Because of stochastic failure of the GET_HOMOLOGUES pipeline to reconstruct some gene family presence/absence profiles, the GET_HOMOLOGUES analysis was run twice, once on a dataset containing *All. vitis sensu stricto* and *All. ampelinum* strains only, and once on a dataset that also included strain Av2, to allow a near-complete coverage of the pan-genome gene families.
